# Another chemolithotrophic metabolism missing in nature—sulfur comproportionation

**DOI:** 10.1101/820597

**Authors:** Jan P. Amend, Heidi S. Aronson, Jennifer Macalady, Douglas E. LaRowe

**Author notes:** Corresponding Author: Jan P. Amend, Department of Earth Sciences, University of Southern California, Zumberge Hall of Science, Los Angeles, CA 90089, USA.

## Abstract

Chemotrophic microorganisms gain energy for cellular functions by catalyzing oxidation-reduction (redox) reactions that are out of equilibrium. Calculations of the Gibbs energy (Δ*G*_*r*_) can identify whether a reaction is thermodynamically favorable, and the accompanying energy yield at the temperature, pressure, and chemical composition in the system of interest. Based on carefully calculated values of Δ*G*_*r*_, we predict a novel microbial metabolism—sulfur comproportionation (3H_2_S + SO_4_^2-^ + 2H^+^ = 4S^0^ + 4H_2_O). We show that at elevated concentrations of sulfide and sulfate in acidic environments over a broad temperature range, this putative metabolism can be exergonic (Δ*G*_*r*_<0), yielding ∼30-50 kJ/mol. We suggest that this may be sufficient energy to support a chemolithotrophic metabolism currently missing in nature. Other versions of this metabolism, to thiosulfate (H_2_S + SO_4_^2-^ = S_2_O_3_^2-^ + H_2_O) and to sulfite (H_2_S + 3SO_4_^2-^ = 4SO_3_^2-^ + 2H^+^), are only moderately exergonic or endergonic even at ideal geochemical conditions. Natural and impacted environments, including sulfidic karst systems, shallow-sea hydrothermal vents, sites of acid mine drainage, and acid-sulfate crater lakes, may be ideal hunting grounds for finding microbial sulfur comproportionators.

## INTRODUCTION

Reaction energetics have been used, at least in part, to posit the existence of several previously undetected microbial metabolisms. The core argument reads that sufficiently exergonic redox reactions can drive cellular functions of hypothetical microorganisms. In a classic example, Broda (1977) calculated standard Gibbs energies to propose that anaerobic ammonia oxidation with nitrate or nitrite (later termed anammox) could fuel certain chemolithotrophs “missing in nature” (1). It took nearly 20 years until microbial catalysis of this process was confirmed in a laboratory wastewater sludge reactor (2), and several more years until it was documented in natural environments with nitrite as the oxidant (3). It is now recognized that the activity of anammox bacteria may account for as much as half of the nitrogen turnover in marine sediments (4).

In another example, anaerobic oxidation of methane (AOM) by marine sediment microorganisms was hypothesized by Barnes and Goldberg (1976) on the strengths of methane and sulfate concentration profiles from the anoxic sediments in the Santa Barbara Basin (California, USA), and on a modest negative free energy change calculated for the *in situ* conditions (5). Other geochemical evidence for AOM in methane-sulfate transition zones of marine sediments followed (6-8). Lipid biomarkers, gene surveys, fluorescence microscopy, and carbon isotopes then confirmed a structured symbiotic relationship between a methane-oxidizing archaeon and a sulfate-reducing bacterium (9-11).

A third example, which puzzled microbiologists for over a century, was the apparent lack in nature of complete ammonia oxidation (comammox). Based again in part on a thermodynamic argument, an organism with this putative metabolism should have growth advantages—but perhaps not kinetic advantages—over incomplete ammonia oxidizers (12). Analogous to the first two scenarios noted above, comammox bacteria were subsequently identified, cultivated, and characterized from an elevated temperature biofilm in an oil exploration well (13) and from a recirculating aquaculture system (14). Their environmental distribution and ecological significance, however, are still largely speculative (15).

Here, we use a thermodynamic approach to predict the existence of another novel microbial metabolism—sulfur comproportionation. Based on geochemical parameters, we then identify a number of natural and impacted target environments where sulfur comproportionation is energetically favorable and where finding microorganisms capable of this metabolism may be possible.

## RESULTS AND DISCUSSION

Sulfur comproportionation can be interpreted either as the anaerobic oxidation of sulfide with sulfate as the electron acceptor, or conversely, as the lithotrophic reduction of sulfate with sulfide as the electron donor. This process can be written to various intermediate oxidation state sulfur compounds, including to elemental sulfur as

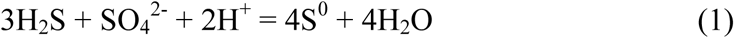

to thiosulfate as

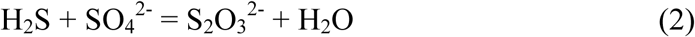

and to sulfite as

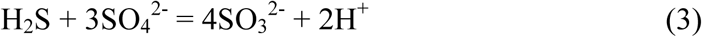

Note that the reverse of Reactions (1), (2), and (3) describes sulfur disproportionation, a confirmed metabolism carried out by, among others, members of the Deltaproteobacteria and Thermodesulfobacteria in several natural environments, including marine sediments, shallow- and deep-sea hydrothermal systems, and alkaline hot springs (16-21). Whether the forward (comproportionation) or reverse (disproportionation) reaction is thermodynamically favorable depends predominantly on the chemical composition of the environment under consideration; temperature and pressure play a far more limited role.

The direction in which a reaction can proceed and the associated energy yield are readily determined with the relation

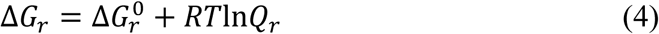

where Δ*G*_*r*_ and 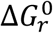 stand for the *overall* and *standard* Gibbs energy of reaction *r*, respectively, *R* and *T* represent the gas constant and temperature (K), respectively, and *Q*_*r*_ denotes the activity product. Values of *Q*_*r*_ can be calculated with the expression

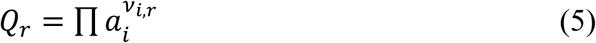

where *a*_*i*_ represents the activity (unitless) of the *i*^th^ species, and *v*_*i,r*_ stands for the corresponding stoichiometric reaction coefficient. The activities of liquid water and pure solids, including S^0^ in Reaction (1), are typically set to unity (1.0), but those of aqueous solutes are computed with the equation

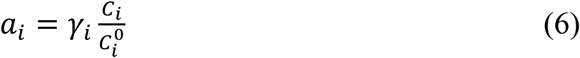

where *γ*_*i*_, *C*_*i*_, and 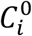 stand for, respectively, the activity coefficient (unitless), concentration (usually in molality, *m*), and standard state concentration (usually 1 *m*). Values of *γ*_*i*_ can be calculated with an expression of the Debye-Hückel equation that takes into account the ionic strength of the solution, and the charge and other properties of the solute species. For recent discussions on this topic, the reader can consult Dick (22), LaRowe and Amend (23), and references therein.

Energy yields for Reaction (1) were calculated at elevated, but geochemically reasonable concentrations of aqueous sulfide and sulfate, and plotted in Fig. 1 as a function of temperature and pH. It can be seen in this figure that Reaction (1) is exergonic (Δ*G*_1_<0) at acidic conditions over the entire temperature range (0-100°C) considered here. It is thermodynamically most favorable, with energy yields exceeding 30 kJ/mol, at pH <≈3 and temperature <≈50°C; energy yields are greater than ∼50 kJ/mol at pH <≈1 and temperature <≈20°C. We should note that the activity of total sulfate (10^−2^) and that of total sulfide (10^−3^) were equilibrium partitioned among HSO_4_^-^ and SO_4_^2-^, and among H_2_S and HS^-^, respectively, at the given temperature and pH. These calculations demonstrate that over the entire range of conditions considered here, 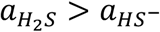, and for the vast majority of the conditions, 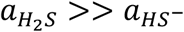. For sulfate, however, values of 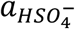 and 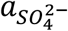 are approximately equal along a curve from pH 1.8 and 0°C to pH 3.0 and 100°C.

**Caption for Figure 1.**
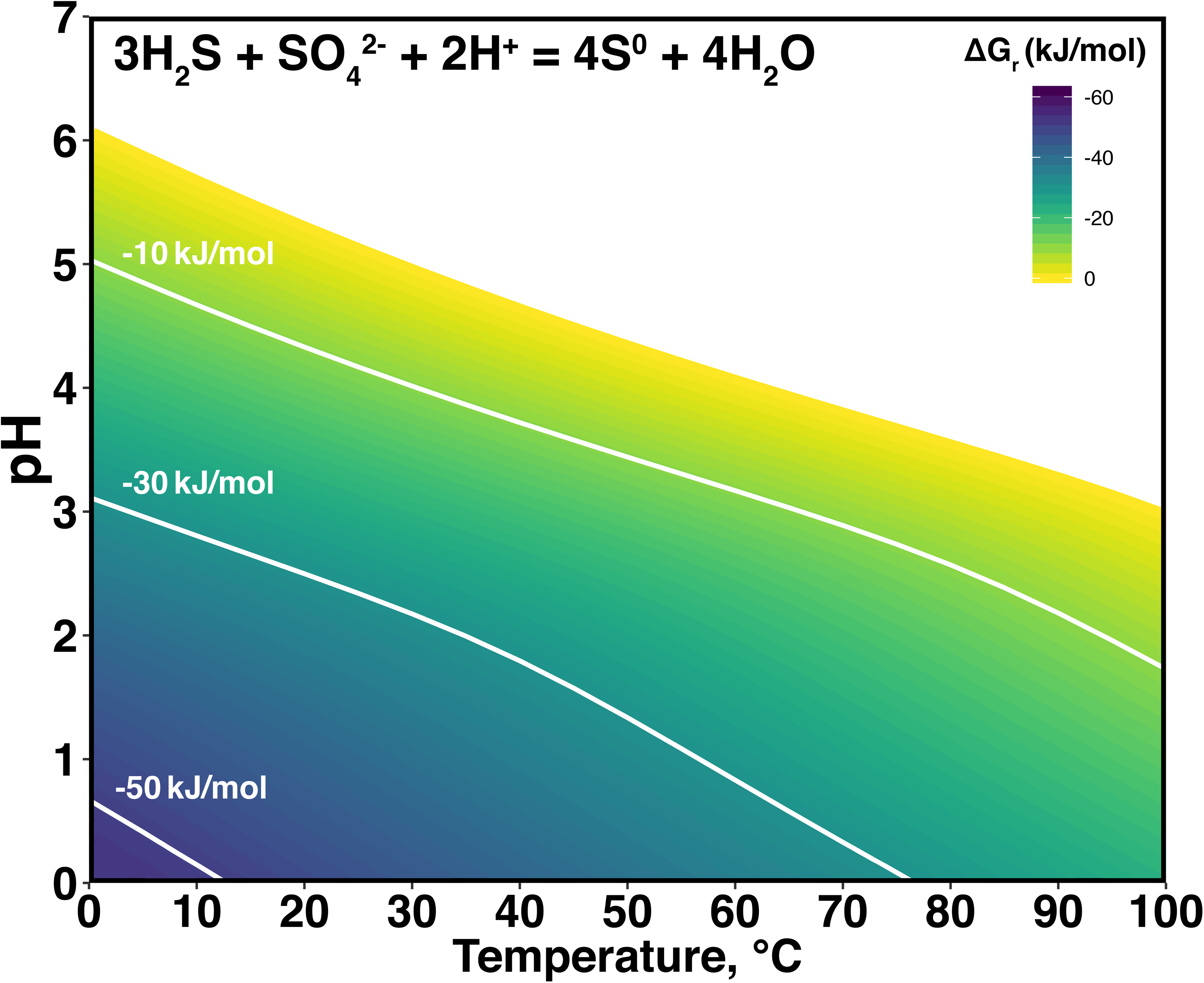
Values of Δ*G*_*r*_ for Reaction (1) calculated with Equation (3) as a function of temperature and pH. Values of 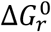 were computed with the SUPCRT92 software package (36). Activities of aqueous H_2_S and SO_4_^2-^ across the temperature and pH space represented here were calculated with equilibrium speciation among H_2_S and HS^-^ given a total sulfide activity of 10^−3^, and among HSO_4_^-^ and SO_4_^2-^ given a total sulfate activity of 10^−2^.

Sulfur comproportionation to thiosulfate (Reaction (2)) can also be exergonic, but only moderately so. Regardless of temperature, it requires high concentrations of sulfate and sulfide and very low concentrations of thiosulfate, but it is not limited to acidic pHs. As an example, values of Δ*G*_2_ at activities of sulfate, sulfide, and thiosulfate equal to 10^−2^, 10^−3^, and 10^−9^, respectively, vary from approximately −9 kJ/mol near 0°C to approximately −15 kJ/mol near 100°C. Sulfur comproportionation to sulfite (Reaction (3)) is an unlikely, perhaps impossible metabolism; energy calculations show that it is endergonic (Δ*G*_3_ >0) even at geochemically optimal conditions (high temperature, alkaline pH, elevated sulfate and sulfide activities, and very low sulfite activities).

By comparison, S^0^ disproportionation can also be exergonic, but at very different geochemical conditions. For example, the energy yield for the reverse of Reaction (1) is ∼96 kJ/mol at 25°C, low levels of SO_4_^2-^ and H_2_S (activities set to 10^−4^ and 10^−6^, respectively), in slightly alkaline aqueous solutions (pH 8). At these conditions and activity of thiosulfate equal 10^−6^, thiosulfate disproportionation represented by the reverse of Reaction (2) yields ∼35 kJ/mol. Finally, at these conditions, activity of sulfite equal 10^−6^, and neutral pH, sulfite disproportionation (reverse of Reaction (3)), yields ∼200 kJ/mol. Recall that all three of these disproportionation reactions have been demonstrated to support the growth of numerous strains of bacteria. We suggest here that if an energy yield as low as ∼35 kJ/mol is sufficient for sulfur disproportionators, then ∼30-50 kJ/mol may similarly suffice to fuel the maintenance and growth of putative sulfur comproportionators.

Many natural and impacted environments exhibit geochemical conditions that are thermodynamically favorable for sulfur comproportionation, and hence may serve as ideal hunting grounds for microorganisms that catalyze this proposed metabolism. As noted, Reaction (3) is endergonic even at the most favorable conditions, but Reaction (2) might be a possibility, perhaps in marine oxygen minimum zones (24, 25). The best conditions for Reaction (1) from an energy standpoint are acidic (pH<3), high sulfate (>50 mM), high sulfide (>1 mM), and low temperature (<20°C); the temperature dependence of the energy yield is rather moderate, however, and Δ*G*_1_ can be reasonably exergonic even at temperatures approaching 100°C. It should be clear that different combinations of these geochemical parameters may also result in a thermodynamic drive for sulfur comproportionation, especially the version represented by Reaction (1).

In cool sulfidic karst systems, formed by the dissolution with sulfuric acid of carbonate rock, aqueous solutions associated with biofilms on the walls (‘snottites’) can be extremely acidic (pH<2.5) and exposed to cave air rich in hydrogen sulfide (∼10-30 ppmv). High value targets include, but are certainly not limited to, the Frasassi cave system in Italy and La Cueva de Villa Luz in Mexico (26, 27). In shallow-sea and surf-zone hydrothermal systems, interfaces between acidic, sulfidic vent fluids and cooler, sulfate-rich seawater may be energy-yielding for sulfur comproportionators. For example, at Vulcano Island (Italy) mixed fluids (∼40-90°C) have been described with pH as low as ∼2, sulfate levels up to ∼60 mM, and sulfide up to ∼0.4 mM (28, 29). At Milos Island (Greece) fluids are lower in sulfate (up to ∼33 mM), but more sulfidic (up to 3 mM) (30, 31). The dissolution and subsequent oxidation of sulfide minerals in acid mine drainage sites can also yield optimum conditions for sulfur comproportionators. For example, at the Richmond Mine (California, USA), temperatures are moderate (30-50°C), pHs can be negative, and measured sulfate concentrations are extremely high (up to 1161 mM) (32, 33). Lastly, the search for sulfur comproportionation may be successful in acid-sulfate crater lakes. There, as for example at Kawah Ijen Volcano (Indonesia), the lake temperature is moderate (37°C), pH is near zero (0.2-0.7), and sulfate levels are high (up to ∼770 mM). Although measured sulfide levels are low (0.02 ppm), H_2_S is likely introduced to the system via fumarolic injections (34, 35).

## CONCLUSIONS

Phototrophs capture and convert the energy from solar radiation to drive all of their cellular processes, but chemotrophs must harness metabolic energy from redox disequilibria. Thermodynamic calculations at the temperature, pressure, and chemical composition of interest enable scientists to propose new combinations of redox couples that may serve as new metabolisms. Here, we demonstrated the potential energy yields for sulfur comproportionation at geochemically reasonable conditions. This approach can be expanded to speculate on other overlooked metabolic strategies, including, but certainly not limited to, nitrogen-redox, metal-redox, and mineral-redox reactions. Such reaction energetics can be used as a hypothesis generator, taken up by geochemists to identify the most likely target environments and by microbiologists to design culturing strategies to find and grow chemotrophs on metabolisms currently missing in nature.

## ACKNOWLEDGMENTS

Funding to JPA and DEL was provided by the Center for Dark Energy Biosphere Investigations (NSF OCE 1431598). Additional support was provided to DEL by the USC Zumberge Fund Individual Grant, the NASA-NSF Origins of Life Ideas Lab program under grant NNN13D466T, and the Alfred P. Sloan Foundation through the Deep Carbon Observatory. HA was supported by the NSF Graduate Research Fellowship Program. This is C-DEBI contribution number XXX.

